# inClust: a general framework for clustering that integrates data from multiple sources

**DOI:** 10.1101/2022.05.27.493706

**Authors:** Lifei Wang, Rui Nie, Zhang Zhang, Weiwei Gu, Shuo Wang, Anqi Wang, Jiang Zhang, Jun Cai

**Author notes:** Correspondence to: Lifei Wang, Jiang Zhang, Jun Cai. These authors contributed equally for this paper (Lifei Wang, Rui Nie).

## Abstract

Clustering is one of the most commonly used methods in single-cell RNA sequencing (scRNA-seq) data analysis and other fields of biology. Traditional clustering methods usually use data from a single source as the input (e.g. scRNA-seq data). However, as the data become more and more complex and contain information from multiple sources, a clustering method that could integrate multiple data is required. Here, we present inClust (integrated clustering), a clustering method that integrates information from multiple sources based on variational autoencoder and vector arithmetic in latent space. inClust perform information integration and clustering jointly, meanwhile it could utilize the labeling information from data as regulation information. It is a flexible framework that can accomplish different tasks under different modes, ranging from supervised to unsupervised. We demonstrate the capability of inClust in the tasks of conditional out-of-distribution generation under supervised mode; label transfer under semi-supervised mode and guided clustering mode; spatial domain identification under unsupervised mode. inClust performs well in all tasks, indicating that it is an excellent general framework for clustering and task-related clustering in the era of multi-omics.

## Introduction

Since the first proposal, the field of single-cell transcriptomics has developed rapidly, and data has accumulated at an unprecedented speed (Chen, et al., 2019). At the same time, the emergence of related new technology, such as spatial transcriptomics, provides a new perspective for understanding basic biological problems (Liao, et al., 2021; Rodriques, et al., 2019; Ståhl, et al., 2016; Vickovic, et al., 2019). However, due to the non-biological differences between data, data of related tissue from different sources couldn’t be directly put together for analysis. For example, the batch effect, that is, the systematic gene expression difference between batches, must be eliminated by batch effect removal methods (e.g.: MNN, CCA) before the integrated downstream analysis is applied to all datasets (Butler, et al., 2018; Haghverdi, et al., 2018). The recently coined term “harmonization” refers to the integration of two or more datasets before downstream analysis(Xu, et al., 2021).

After harmonization, several downstream analyses could be conducted. The most preliminary and important downstream step is clustering, which is the basis for cell type identification and further analysis (Kiselev, et al., 2019). Because harmonization and downstream analysis are closely related, it is best to implement them jointly. There are several methods to run harmonization and clustering simultaneously, such as harmony and DESC (Korsunsky, et al., 2019; Li, et al., 2020). However, as unsupervised methods, they couldn’t make effective use of labeling information, which usually exists in the dataset called the reference. Then, the semi-supervised methods that could use labeling information in reference datasets are developed, such as scAdapt and scANVI(Xu, et al., 2021; Zhou, et al., 2021). Those methods co-cluster labeled data in reference dataset and unlabeled data in target dataset, then transfer labels from reference dataset to target datasets.

Deep generative models are widely used in single cell data analysis(Gayoso, et al., 2022). The methods based on the deep generative model could perform harmonization (scVI) or clustering (scVAE) alone(Grønbech, et al., 2020; Lopez, et al., 2018), or run harmonization and semi-supervsied cell type annotation jointly (scANVI). In addition, these types of models could accomplish various tasks, such as multi-modal omics data analysis (total VI)(Gayoso, et al., 2021), imputation of missing genes in spatial transcriptomics from scRNA-seq data (gimVI)(Lopez, et al., 2019), or prediction of perturbation responses in conditional out-of-distribution generation problem (scGen and trVAE)(Lotfollahi, et al., 2020; Lotfollahi, et al., 2019). Furthermore, the new development of a transfer learning framework, called single-cell architectural surgery (scArches), allows nodes with parameters and datasets (query) to be added to existing architecture and datasets (reference) with minimal modification to the original deep generative model (architecture surgery)(Lotfollahi, et al., 2022). Applying scArches to deep conditional models (e.g.: trVAE, scVI, scANVI, totalVI) greatly enhances the capabilities of those models.

However, these deep generative models still have some limitations and deficiencies. Most models do not make full use of the information stored in the data, especially the cell type information, which is effectively used only by a few methods (e.g.: scANVI). In solving of out-of-distribution generation problem, cell type information could provide another layer of constraint, but existing methods couldn’t combine this information (e.g.: scGen and trVAE). Meanwhile, different models are implemented for different tasks, such as clustering, annotation of cell-type, or prediction of the perturbation responses. There is not a general framework providing solutions for all those tasks.

Here, we present inClust (integrated clustering), a general clustering method that integrates information from multiple sources. inClust is based on conditional variational autoencoder (cVAE)(Kingma and Welling, 2013), a deep generative model. In the canonical example of conditional variational autoencoder, it encodes scRNA-seq data and batch information jointly in the inference process, then injects the batch information back into the model in the generative process(Gayoso, et al., 2022). Different from that, inClust encode scRNA-seq data and batch information (or other covariates and auxiliary information) into latent space, respectively. So, the influence of the batch and other covariates is explicitly eliminated by vector arithmetic in latent space, unlike implicit removal in the canonical example of conditional variational autoencoder. The latent representations after removing the influence of batch and other covariates are used as the starting point for clustering (without label) or classification (with label). Therefore, the reconstruction part and the additional functional part (covariate effect removal and clustering) are relatively modular, running in parallel but connected with each other (Fig 1A). inClust provides a general and flexible framework, which can be applied to various tasks with different modes. We apply inClust in four tasks: prediction perturbance responses in conditional out-of-distribution generation problems, transferring labels between datasets, transferring labels between partially overlapped datasets, identification of spatial domains in spatial transcriptomics; with supervised, semi-supervised, guided, the unsupervised mode corresponding to each task (Fig 1B-E). The different mode is due to the difference in the proportion of the labeled data in the overall dataset and the purpose of the task. We have proved that inClust is excellent in all tasks.

**Figure 1.**
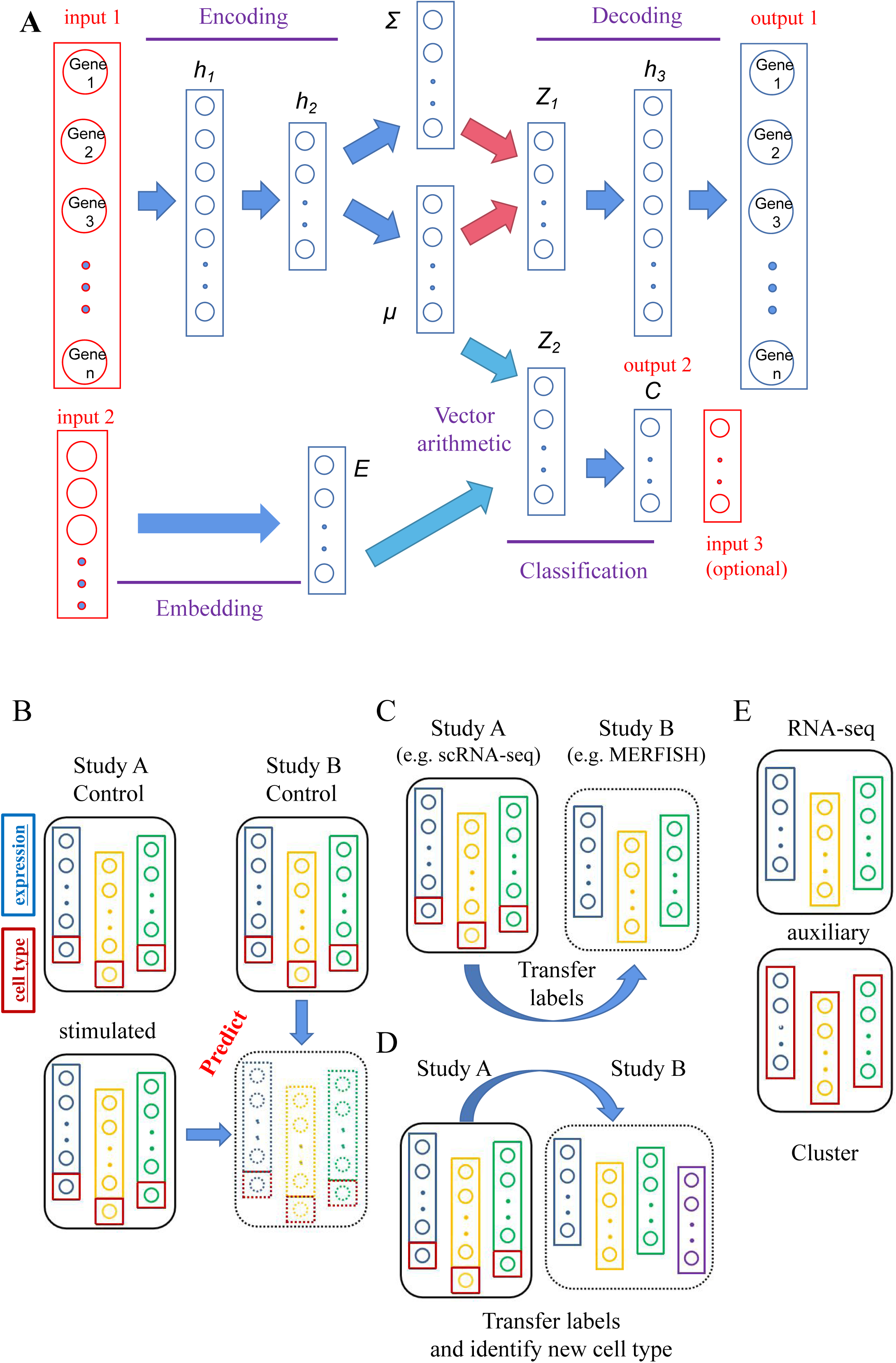
Architecture of inClust and its wide application in field of transcriptome A. The architecture of inClust. The inClust is based on variational auto-encoder, so its backbone contains an encoder, sampling part, and a decoder as VAE. In inClust, both the encoder and decoder are 2 layer neural networks with a non-linear function as the activation function. The sampling part generates parameters (mean and standard deviation) for posterior distribution, which are inputs for reparameterization tricks. In addition to the VAE backbone, inClust has built-in three additional parts, namely, an embedding layer embedded auxiliary information (e.g.: covariates such as batch, donor…) into the latent space, the vector arithmetic part performs mixing of auxiliary information and the input in the latent space, a classifier group cells into clusters. B-E. Four kinds of applications under different modes. Expressions are represented by colored rectangles, and corresponding cell types are represented by red squares. B. The conditional out-of-distribution generation problem. Two scRNA-seq studies, study A and study B, are involved. Control state and stimulated state in study A, control state in study B are known (solid black frame), and the goal is to produce stimulated state in study B (dotted black frame). C. Semi-supervised label transferring. The task is to transfer labels from reference dataset (study A, solid black frame) to target dataset (study B, dotted black frame). D. Guided cell type annotation between partially overlapped datasets. The task is to transfer labels from reference dataset (study A, solid black frame) to multiple target datasets (study B, dotted black frame) and identify a new cell type in the target datasets (purple rectangle in dotted black frame) simultaneously. E. integrated unsupervised clustering. The task is to cluster based on the integrated information from transcriptome and auxiliary information.

## Methods

### Data sets and preprocessing

The scRNA-seq dataset of PBMC was from Zheng et al. and Kang et al.(Kang, et al., 2018; Zheng, et al., 2017). Then, according to the preprocessing method of Lotfollahi et al.(Lotfollahi, et al., 2019), we obtained 7000 genes and 17516 cells. All data was already normalizedand log scale before being used.

The scRNA-seq dataset and spatial transcriptomics dataset of mouse brain were obtained from Gene Expression Omnibus (GSE113576) and Dryad repositories, respectively(Moffitt, et al., 2018). Then, according to the method of Zhou et al.(Zhou, et al., 2021), we obtained 154 genes and 104025 samples.

The scRNA-seq datasets of human heart were obtained from Hua et al. and Litviňuková et al.(Hua, et al., 2020; Litviňuková, et al., 2020). We selected Donor 1, 2, and H3 for analysis. Then, according to the method of Chen et al(Chen, et al., 2022), we obtained 43,878 genes. We selected variable genes with Seurat V4.1(Hao, et al., 2021). After that, we obtained 5000 genes and 83464 cells.

The spatial gene expression and histology image data of the mouse posterior brain were downloaded from https://drive.google.com/drive/folders/1zten54vkjorp26T4iD0ApQGa9ut5eY42?usp=sharing(Hu, et al., 2021), we obtained 18474 genes and 3353 samples. All data was already normalized and log scale before being used.

### Architecture of inClust (Fig 1A)

#### Input

inClust take 3 inputs, input 1 is the scRNA-seq data (real-valued vector) and input 2 is the auxiliary information (real-valued vector or one-hot vector) such as covariates (e.g. batch). Input 3 is the label information (one-hot vector), which is optional.

#### Encoder

The encoder is a three-layer neural network with decreasing dimensions (input1, h1, h2). Each layer uses a non-linear function as the activation function.

#### Latent sampling layer

Both parameters mean (μ) and standard deviation (Σ) are estimated from h2, using a neural network without activation function. The reparameterization trick was used for sampling latent variables Z1.

#### Embedding layer

The embedding layer embeds the auxiliary information (input2) into the latent space as a real-valued vector (E). For example, the unwanted covariates (e.g.: batch, donor) or auxiliary information that are represented by a one-hot vector or real-valued vector are embedded into a real-valued vector in latent space.

#### Vector arithmetic layer

The vector arithmetic is performed in the latent space. The estimated mean (μ) would substrate (or add) the embedding vector E. The resulting vector Z2 retains the real biological information after removing the unwanted covariates or mixing the auxiliary information E.

#### Classifier

The real-valued vector Z2 will pass through a neural network with softmax as the activation function. The output of the classifier is the output2.

#### Decoder

The decoder is a three-layer neural network with increasing dimensions (z1, h3, output1). Each layer uses a non-linear function as the activation function.

## Results

### inClust could predict the perturbation in conditional out-of-distribution generation

Conditional out-of-distribution generation is very common in the study of scRNA-seq(Lotfollahi, et al., 2020). For example, there are 2 studies on the human peripheral blood mononuclear cells (PBMCs). In study A, both control and stimulation conditions (stimulation with interferon IFN-β) are known(Kang, et al., 2018); In study B, only the situation of the control group was known(Zheng, et al., 2017). The purpose of conditional out-of-distribution generation is to predict the perturbation caused by the IFN-β in study B based on the results of study A (control and stimulation). In other words, conditional out-of-distribution generation is to extend the results from one study to others. Methods to solve these problems have been put forward, but most of them do not make good use of all the information. For example, scGen and trVAE, two deep generative models, do not fully use the cell type information existing in the original data (Lotfollahi, et al., 2020; Lotfollahi, et al., 2019).

inClust makes full use of the labeling information in the dataset and provides a solution to such problems in supervised mode. It could transfer finding across multiple studies where scRNA-seq expression data, covariate information (e.g.: batch, donor, technology), and cell types are known. Though simultaneously encoding (embed) the expression data and covariate information (e.g.: batch, condition) into latent space, inClust explicitly eliminates the influence of the covariates in latent space by vector arithmetic and then clustering the samples with unified representations into biologically consistent groups under the supervision of known cell types (Fig 2A). So, the stimulated state in study B could be generated by adding the unified representation of the stimulated state in study A to the latent representation of covariate of stimulated state in study B (Fig 2B, upper: cross batch); or by adding the unified representation of control state in study B and the latent representation of covariate of stimulated state in study B (Fig 2B, lower: cross condition).

**Figure 2.**
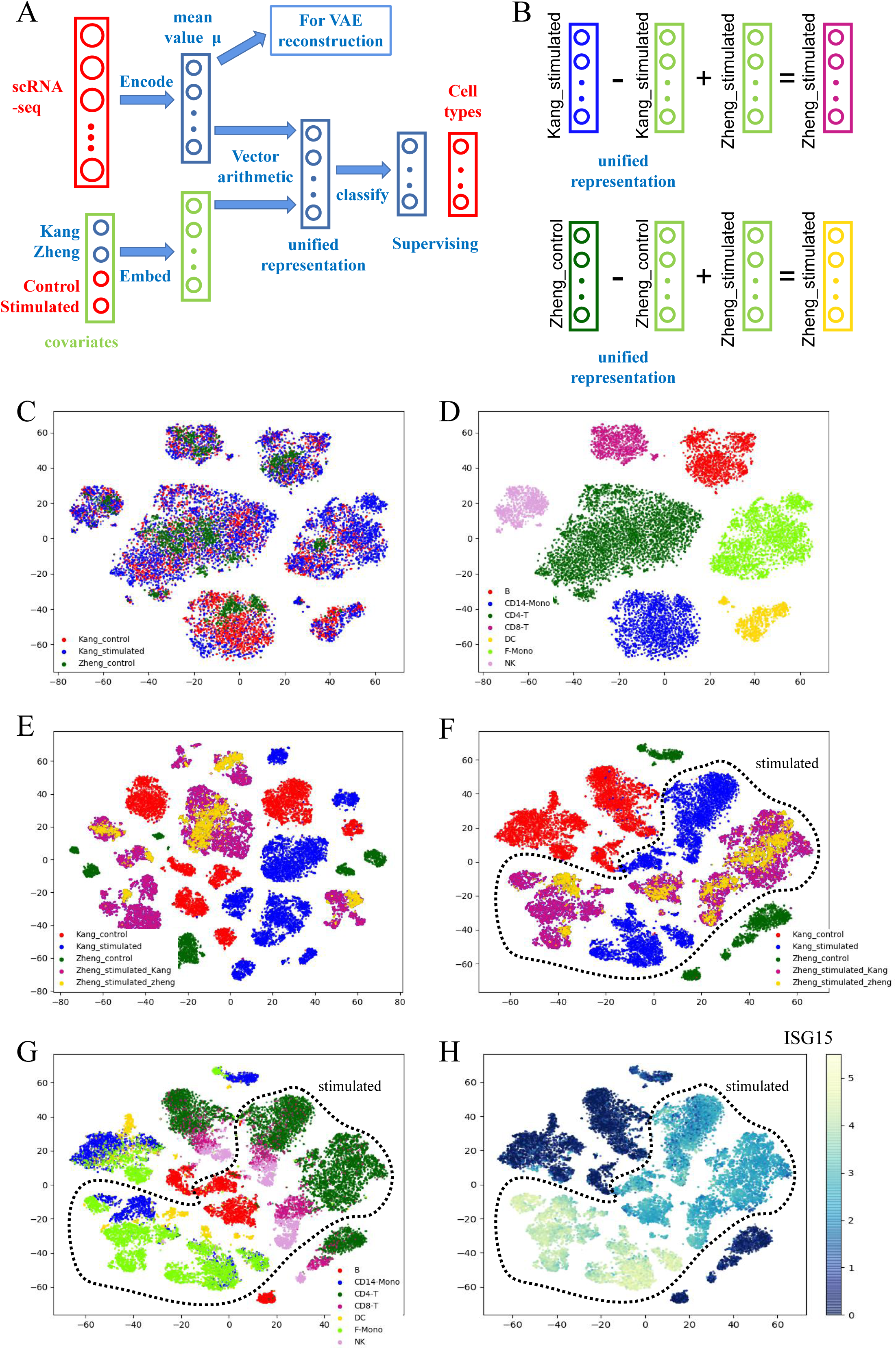
inClust predicts the perturbation response in task of conditional out-of-distribution generation under supervised mode A. Diagram of inClust with supervised mode. Two covariates (study and condition) are represented as two-hot vectors with dimensions of four. The cell type label for each sample is used as the supervision information to constrain the model. B. vector arithmetic in latent space to construct stimulated state latent mean value representation in study B (Zheng_stimulated). Two ways to construct stimulated state latent mean vector, in both ways the unified latent representation is generated by first substrate latent representation of the corresponding covariate from the latent representation of expression data, then add the latent representation of the stimulated state in study B (Zheng_stimulated). Kang: study A; Zheng: Study B. C. The t-SNE plot of the latent unified representation by inClust is colored by study and condition. D. The t-SNE plot of the latent unified representation by inClust is colored by cell type. E. The t-SNE plot of the latent mean value representation by inClust is colored by study and condition after the construction of stimulated state latent mean value representation in study B. The two kinds of constructed stimulated state latent mean value representation in study B are plotted as purple and gold in the plot respectively. F. The t-SNE plot of the expression data (the top 50 PCs) is colored by study and condition. The generated expression data from stimulated state latent mean value representation in study B are plotted as purple and gold in the plot respectively. The stimulated state is contained in the dashed line. G. The t-SNE plot of the expression data (the top 50 PCs) is colored by cell types. The stimulated state is contained in the dashed line. H. The expression value of ISG15 in cells among studies and conditions. The stimulated state is contained in the dashed line.

We apply inClust in the above-mentioned human PBMCs data. The results show that, after covariates (studies and conditions) removal, samples across studies are under the unified representation in latent space, where samples from different studies and conditions are mixed together (Fig 2C), and clustered in the biologically meaningful cell types (Fig 2D). Before removing the covariates, the samples are also separated by studies and conditions, except for the cell types (Fig S1A, B). In latent space, the stimulated state in study B could be constructed from the stimulated state in study A through cross-batch correction (purple in Fig 2E), or from control state in study B through cross-condition correction (gold in Fig 2E). Both constructed stimulated states in study B are clustering together in latent space (Fig 2E, Fig S1C), following our assumption (Fig 2B). Furthermore, as a generative model, inClust could generate the expression data from the vectors in latent space. The expression data generated from latent representation in constructed stimulated state are also clustering together like that in latent space (purple and gold in Fig 2F). In the plot, the generated stimulated state in study B and the real stimulated state in study A are located in the center of the figure, while the control states are located in the periphery (Fig 2F, G). As in previous research (Lotfollahi, et al., 2019), the expression of three different types of marker genes was visualized. The expression of perturbation response specific genes is concentrated in stimulated cells (ISG15: Fig 2H, IFI6: Fig S1D, IFIT1: Fig S1E); the expression of cell type-specific perturbation response genes is restricted to parts of stimulated cells (CXCL10 in monocytes and DC: Fig S1F); the expression of cell-type-specific markers are expressed in specific cells with both control state and stimulated state (CD79A for B cells: Fig S1G, CCL5 for NK and CD8 T cells: Fig S1H). All results show that the inClust performs well in predicting the perturbation in conditional out-of-distribution generation.

### inClust could transfer labels between scRNA-seq data and spatial transcriptomics data

With the accumulation of more and more data, a large reference atlas of scRNA-seq has been constructed(Regev, et al., 2017). The cells in the newly synthesized dataset could be annotated by labels transferring from a reference dataset with a similar research target (e.g.: system, organ). It is necessary to transfer labels between studies, especially in the era of spatial transcriptomics. The spatial location of cells could provide important information for solving many basic problems in biology. Recently, several methods have been developed for spatial transcriptomics(Liao, et al., 2021), and many computational tools have been invented to label the spatial transcriptomics data based on scRNA-seq reference (RCTD, Cell2location)(Cable, et al., 2022; Kleshchevnikov, et al., 2022). MERFISH is a novel method that measures spatial position and gene expression simultaneously(Moffitt, et al., 2018). Cell types in MERFISH data could be identified based on scRNA-seq data(Zhou, et al., 2021).

inClust also provides a solution for label transferring through a semi-supervised manner. First, the inClust harmonizes different data, generating unified latent representation across datasets. Then, for samples from reference dataset, the known cell type would be used as supervision information to lead the clustering process; and the samples from target dataset would presumably co-cluster with reference dataset according to the underlying biologically meaning (cell types) (Fig 3A). Therefore, labels can be transferred from the reference dataset to the target dataset within the same cluster.

**Figure 3.**
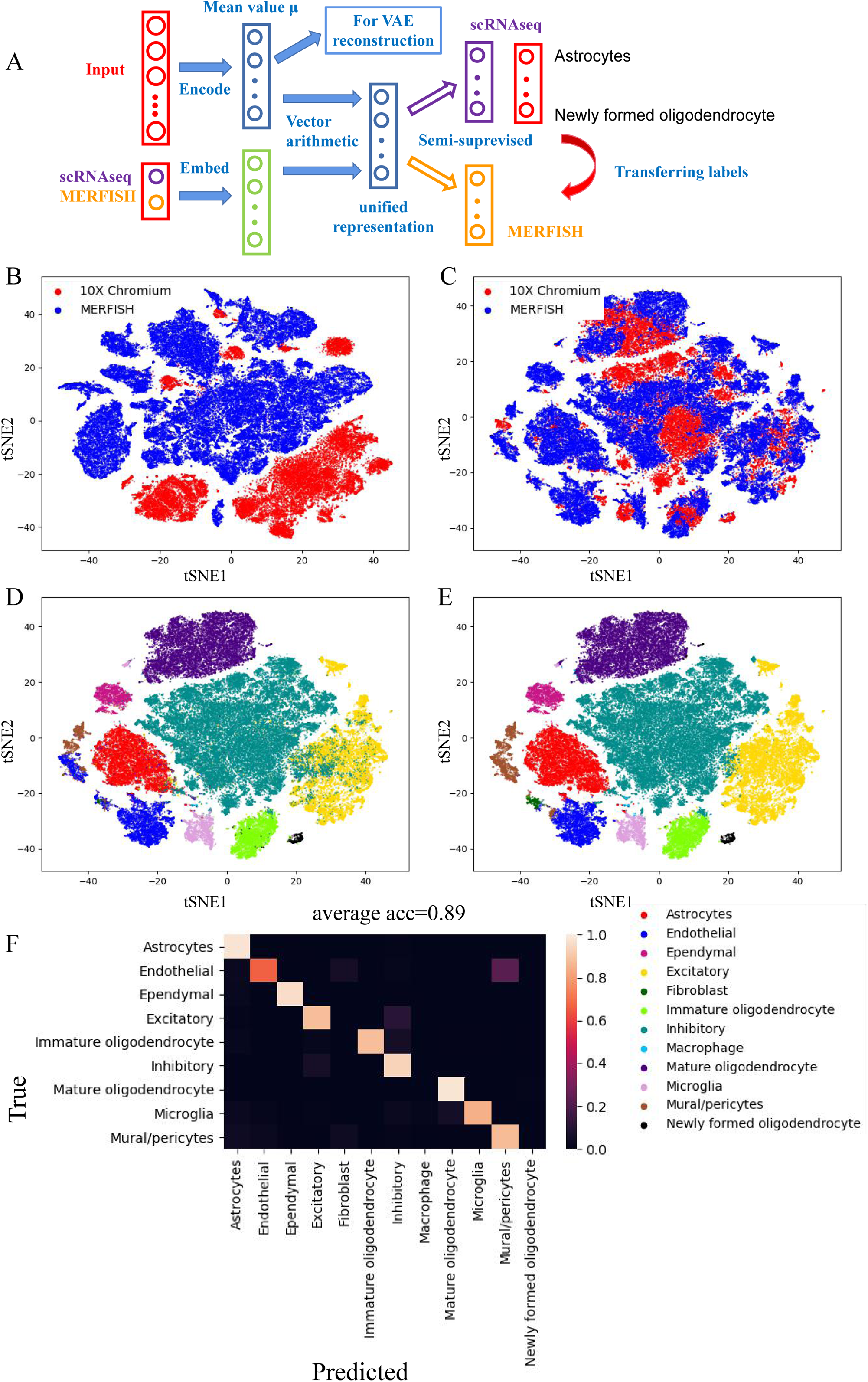
inClust transfer labels between scRNA-seq data (10x Chromium) and spatial transcriptomics data (MERFISH) under semi-supervised mode. A. Diagram of inClust under semi-supervised mode. Two studies (scRNA-seq and MERFISH) are represented as one-hot vectors with dimensions of two. Only the cell type label for scRNA-seq is used as the supervision information to train the model. B. The t-SNE plot of the pre-processed scRNA-seq data and MERFISH data is colored by batch. C. The t-SNE plot of the latent unified representation for scRNA-seq data and MERFISH data in inClust colored by batch. D. The t-SNE plot of the latent unified representation for scRNA-seq data and MERFISH data in inClust colored by cell types. E. The t-SNE plot of the latent unified representation for scRNA-seq data and MERFISH data in inClust colored by predicted cell types. F. Heatmap for the confusion matrix of inClust with average accuracy above.

We test inClust’s capability for data harmonization and label transferring by scRNA-seq data and MERFISH data from the hypothalamic preoptic region(Zhou, et al., 2021). Before data harmonization, cells were grouped according to their sources, in which the cells from scRNA-seq and cells from MERFISH were not mixed (Fig 3B). After data harmonization by inClust, cells from scRNA-seq and MERFISH were mixed together in the latent space (Fig 3C). And cells were clustered into biologically related groups, in which the cells from the same cell types were grouped together (Fig 3D). In the experiments, only cells from scRNA-seq dataset had labels (Fig S2), so the labels were transferred from the scRNA-seq data to MERFISH data in each corresponding group (Fig 3E). Comparing the predicted cell types with the real cell types in the MERFISH dataset, the inClust achieved an average accuracy of 0.89 on nine cell types, which was higher than that reported by scAdapt (Fig 3F).

### inClust could transfer labels between batches and donors, meanwhile identify new cell types in target datasets

Traditional semi-supervised methods usually operate in a one-to-one source-target correspondence at the dataset level and cell type level(Zhou, et al., 2021). However, in many cases, there may be multiple target datasets, and covariates exist between them (e.g.: donor). Meanwhile, the cell type composition between reference dataset and target dataset may partially overlap(Xu, et al., 2021). For example, the target dataset may contain a new cell type. Therefore, not all cells in the target dataset should be assigned labels in the reference dataset.

inClust under guided clustering mode could provide a solution for these situations. In the guided clustering mode, the number of clusters is set to be larger than the number of cell types in the reference dataset. Therefore, the clusters could be divided into semi-supervised clusters (cell type in reference) and unsupervised clusters. In operation, inClust firstly harmonizes multiple datasets with different levels of covariates at once. Then, the semi-supervised clusters are guided by reference dataset, and unsupervised clusters are formed unsupervised. Compared to the semi-supervised mode, the reference dataset merely acts as guiding parts of the clustering process. If target datasets contain cell types that do not exist in the reference dataset, the extra-cluster would account for the new cell types (Fig4 A).

**Figure 4.**
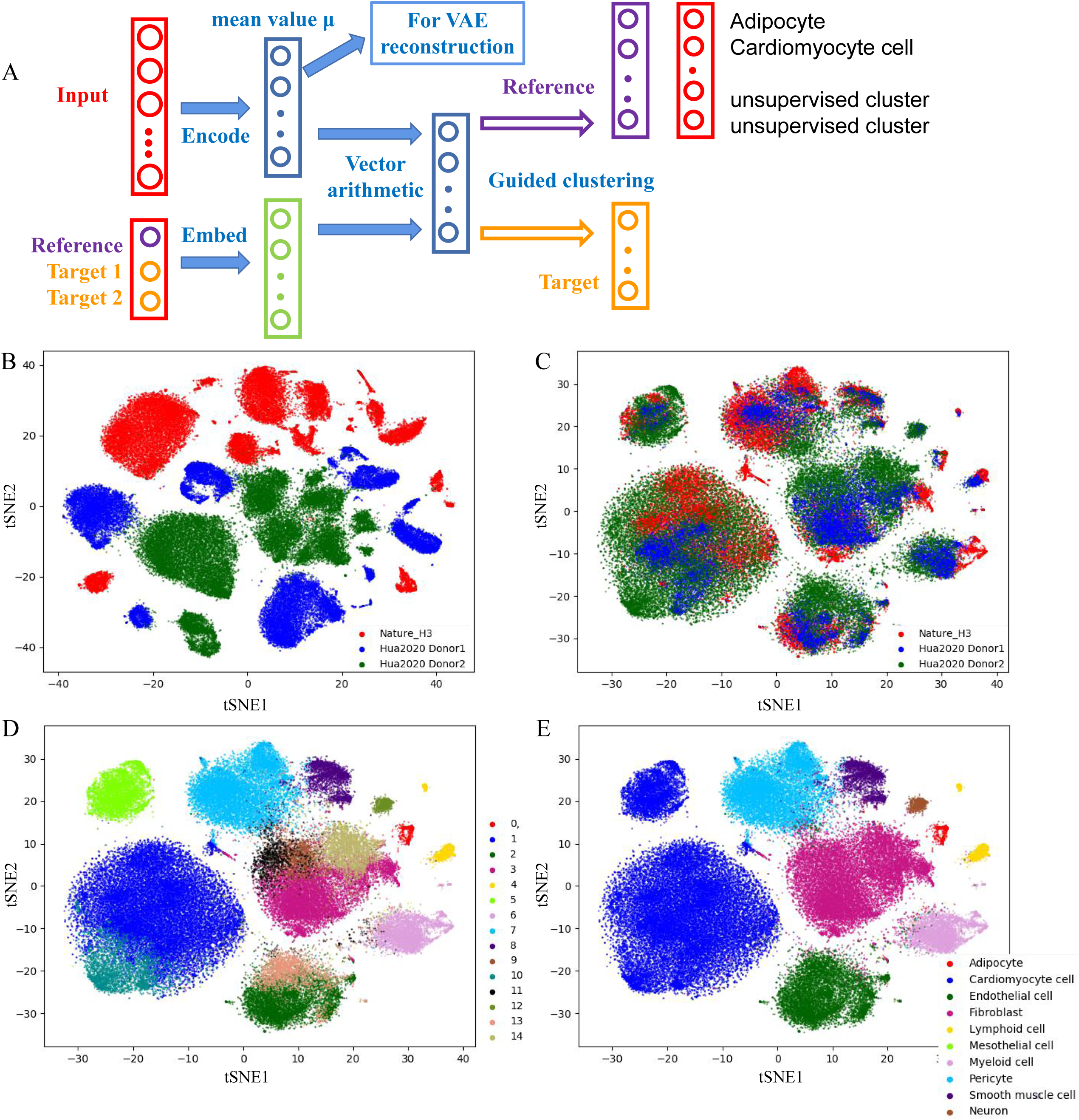
inClust transfers labels between multiple datasets and identifies new cell types in target datasets under guided-clustering mode. A. Diagram of inClust under guide-clustering mode. The covariate (studies and donors) is represented as a one-hot vector with dimensions of three. The number of clusters is set to be greater than the number of cell types in the reference dataset. Therefore, clusters could be divided into semi-supervised clusters (cell type in reference) and unsupervised clusters. The cell type label in the reference dataset is used as the supervision information to constrain the model. The unsupervised clusters could account for new cell types in the target dataset. B. The t-SNE plot of the heart scRNA-seq data is colored by the donor. C. The t-SNE plot of the latent unified representation for heart scRNA-seq data in inClust is colored by donor. D. The t-SNE plot of the latent unified representation for the heart scRNA-seq data in inClust is colored by clusters. E. The t-SNE plot of the latent unified representation for heart scRNA-seq data in inClust is colored by predicted cell types.

To test our model, we used the scRNA-seq dataset of the human heart from 2 studies. The reference dataset contains 8 cell types (‘Adipocyte’, ‘Cardiomyocyte cell’, ‘Endothelial cell’, Fibroblast’, ‘Lymphoid cell’, ‘Myeloid cell’, ‘Pericyte’, ‘Smooth muscle cell’) with labeling information from one donor (Litviňuková, et al., 2020). And the ‘Cardiomyocyte cell’ could be further divided into ‘Atrial cardiomyocyte cell’ and ‘Ventricular cardiomyocyte cell’. The two target datasets are from different donors, and contain ‘neuron’ that is not part of the reference dataset(Hua, et al., 2020). As shown in Fig 4B, there is an obvious batch effect among the three datasets, cells are separated by batches and donors in the t-SNE plot. In contrast, after applying inClust, the batch effect is effectively removed, the cells from different batches and donors are well mixed in the t-SNE (Fig 4C). We set the number of clusters to 15, which is larger than the cell type in the reference dataset, as the clustering results shown (Fig 4D). For the semi-supervised clusters (clusters 0-8), the annotation of the cluster could be easily accomplished by transferring labels from reference datasets (Fig S3A). For the rest unsupervised clusters, the annotation could be achieved by identifying the corresponding marker genes (Fig S3B). For example, the cluster 9 could be labeled as Fibroblast according to marker genes C3, MEG3, and ACSM3(Friscic, et al., 2021; Koenig, et al., 2022; Piccoli, et al., 2017); the cluster 10 could be labeled as Cardiomyocyte cell according to marker genes SPEG, ACTA1 and MYL3(Campbell, et al., 2021; DeLaughter, et al., 2016; Koenig, et al., 2022); the cluster 11 could be labeled as Fibroblast according to marker genes ACSM3(Koenig, et al., 2022); the cluster 12 could be labeled as Neuron according to the marker genes NRXN1 and NRXN3(Litviňuková, et al., 2020); the cluster 13 could be labeled as Endothelial cell according to the marker genes VWF and PECAM1(Litviňuková, et al., 2020); the cluster 14 could be labeled as Fibroblast according to the marker genes FBLN1, FBLN2 and CFH(Litviňuková, et al., 2020; Muhl, et al., 2020). A new cell type ‘Neuron’ has been identified in unsupervised clusters. All results show that the inClust could transfer labels from reference dataset and identify new cell types in the target dataset.

### inClust could identify spatial domains in spatial transcriptomics

As mentioned above, the spatial transcriptomic could measure transcriptomic and spatial location simultaneously. Usually, the spatial transcriptomic is conducted on a 2-D plane. The key step in spatial transcriptomic is to divide the plane into meaningful parts, which is called spatial domain. The primitive way to define spatial domain is to use clusters generated by clustering algorithms (e.g.: K-means and Louvain’s method) only based on gene expression data. But besides gene expression data, a lot of extra information, such as histology, could be used to define the spatial domain. The trend of integrating gene expression data and additional information to define spatial domain is becoming more and more obvious(Hu, et al., 2021; Pham, et al., 2020).

inClust is an integration framework for clustering. It provides a solution for identifying spatial domains by integrating information from multiple sources. As shown in diagram (Fig 5A), inClust takes gene expression data and auxiliary information (RGB value of histological image) as input, then encodes (embeds) this information into the latent space, and integrates information from both parties by vector arithmetic in the latent space. Because the task is to integrate multiple data rather than remove unnecessary covariates, addition is used instead of the substation in other examples. Finally, the clustering is performed on the vectors that integrate information from multiple sources.

**Figure 5.**
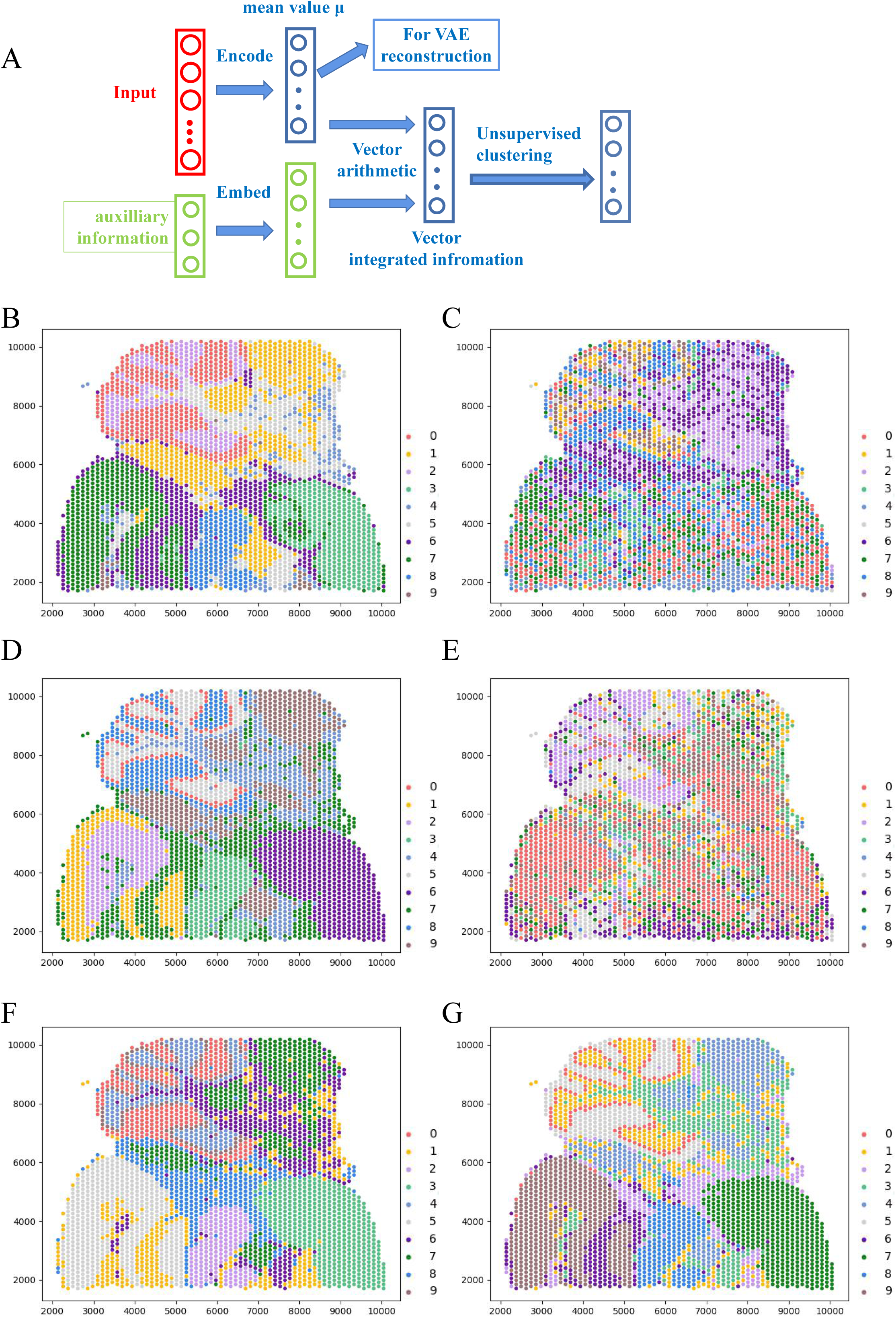
inClust identifies spatial domains in spatial transcriptomics by integrating gene expression data and histological information under unsupervised mode. A. Diagram of inClust under unsupervised mode. The auxiliary information (RGB in histological image) is represented as real-valued vector with dimension of three. The integration step takes place in the vector arithmetic (addition) in latent space, resulting in information integration vector. Then, clustering is carried out by using the information integration vector. B. Spatial domains of the cortex region detected by K-means based on gene expression data alone. C. Spatial domains of the cortex region detected by K-means based on smoothed RGB values of histological image. D. Spatial domains of the cortex region detected by K-means based on latent representation of gene expression data in inClust. E. Spatial domains of the cortex region detected by K-means based on latent representation of smoothed RGB values of histological image in inClust. F. Spatial domains of the cortex region detected by K-means based on latent representation after data integration by vector addition in inClust. G. Spatial domains of the cortex region detected by inClust.

We test inClust’s capability to integrate unsupervised clustering by 10x Visium dataset generated from mouse posterior brain and its associated histological image. As in the previous study, the first 50 principal components of 10x Visium dataset were used as one input for clustering. Meanwhile, the smoothed RGB values (average value of 50× 50 pixel squares around the target pixel, Fig S4 A, B, C) are used as the auxiliary information for another input (Hu, et al., 2021). Before data integration, we use K-means to cluster the two inputs separately. The clustering results are plotted (Fig 5B, C). Then, we use inClust for integrated clustering, and the two inputs are encoded into latent space respectively. We also use the K-mean to cluster the two inputs in latent space, respectively. The spatial domain in latent space seems to be more contiguous than that in original space, especially the histological data (Fig 5D, E). Then, the K-means results of the integrated vector in latent space (Fig 5F) and the cluster output of inClust are plotted (Fig 5G). The plot shows that the results generated by those two methods are similar and seem more contiguous than transformed 10x Visium alone (Fig 5D). Those results demonstrate the inClust’s ability for integrated unsupervised clustering.

## Discussion

In summary, we have proposed a deep generative based model inClust, which could perform information integration and clustering jointly. In addition, label information of cell type in the dataset could be effectively used in inClust. According to the object of the task and the extent to which label information is covered in the overall dataset, inClust could operate under a range of modes, from totally supervised, to semi-supervised and guided clustering, finally to totally unsupervised clustering. One by one, we tested inClust under each mode on associated tasks. First, under the supervised mode, inClust predicts IFN-β response in human PBMC by generalizing IFN-β response across studies. To achieve this goal, inClust integrates gene expression data, study and condition information, and cell type labels in all datasets. Secondly, under the semi-supervised mode, inClust transfers cell type labels from scRNA-seq data to spatial transcriptomics data (MERFISH) in the hypothalamic preoptic region with high accuracy. Thirdly, under guided-clustering mode, inClust transfers cell type labels from reference dataset of the human heart in one study to two target datasets of different donors in another study. At the same time, inClust identified new cell types that only exist in target dataset. Finally, under the unsupervised mode, inClust identified the spatial domains in spatial transcriptomics from mouse posterior brain by integrating gene expression data and histological information. Generally speaking, inClust performs well in every task mentioned above, which proves its ability as a general framework to solve cluster-related work that needs to integrate data from multiple sources.

Deep generative models are widely used in data analysis of scRNA-seq(Gayoso, et al., 2022). Conditional variational autoencoder, one of the deep generative models, is often used as the backbone of the computational model for single cell transcriptome analysis. These computational models (e.g.: scVI, scANVI, and trVAE) could complete tasks such as batch effect removal, cell type annotation, and conditional out-of-distribution generation. The structure of this conditional variational autoencoder is canonical: the scRNA-seq data and covariate conditional information are jointly encoded in the same neural networks during the inference process, to regress out the effect of covariate conditional information; then covariate conditional information is injected back to reconstruct the sample during the generative process. inClust use an alternative way to integrate information: the gene expression data and covariate conditional information is encoded (embed) separately into latent space with different neural networks; then vector arithmetic in latent space is used for information integration. which has proved effective in deep generative model (Lotfollahi, et al., 2019). Furthermore, inClust could flexibly and conveniently incorporate label information. As a result, inClust could complete a series of tasks under different modes, ranging from totally supervised, to semi-supervised and guided clustering, finally to totally unsupervised clustering. In this sense, inClust can be used as a general framework to complete various tasks encountered in scRNA-seq analysis, which were initially completed by several different models.

## Competing financial interestsn

The authors declare no competing financial interests.

## Figure legend

**Figure S1.**
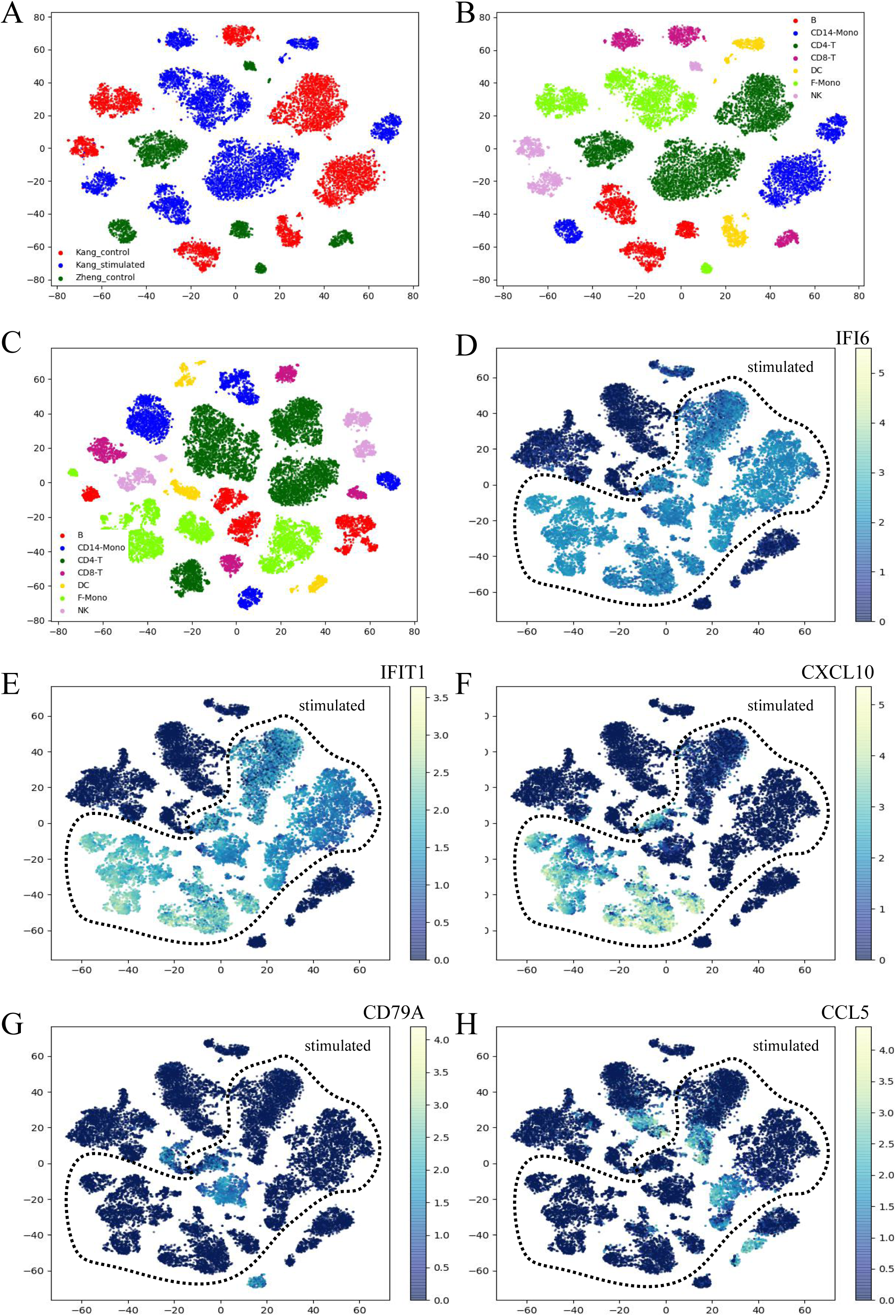
results of conditional out-of-distribution generation task A. The t-SNE plot of the latent mean value representation in inClust is colored by study and condition. Kang: study A; Zheng: Study B. B. The t-SNE plot of the latent mean value representation in inClust colored by cell types. C. The t-SNE plot of the latent mean value representation in inClust colored by cell types after the construction of stimulated state latent mean value representation in study B. D. The expression value of IFI6 in cells among studies and conditions. The stimulated state is contained in the dashed line. E. The expression value of IFIT1 in cells among studies and conditions. The stimulated state is contained in the dashed line. F. The expression value of CXCL10 in cells among studies and conditions. The stimulated state is contained in the dashed line. G. The expression value of CD79A in cells among studies and conditions. The stimulated state is contained in the dashed line. H. The expression value of CCL5 in cell among studies and conditions. The stimulated state is contained in the dashed line.

**Figure S2.**
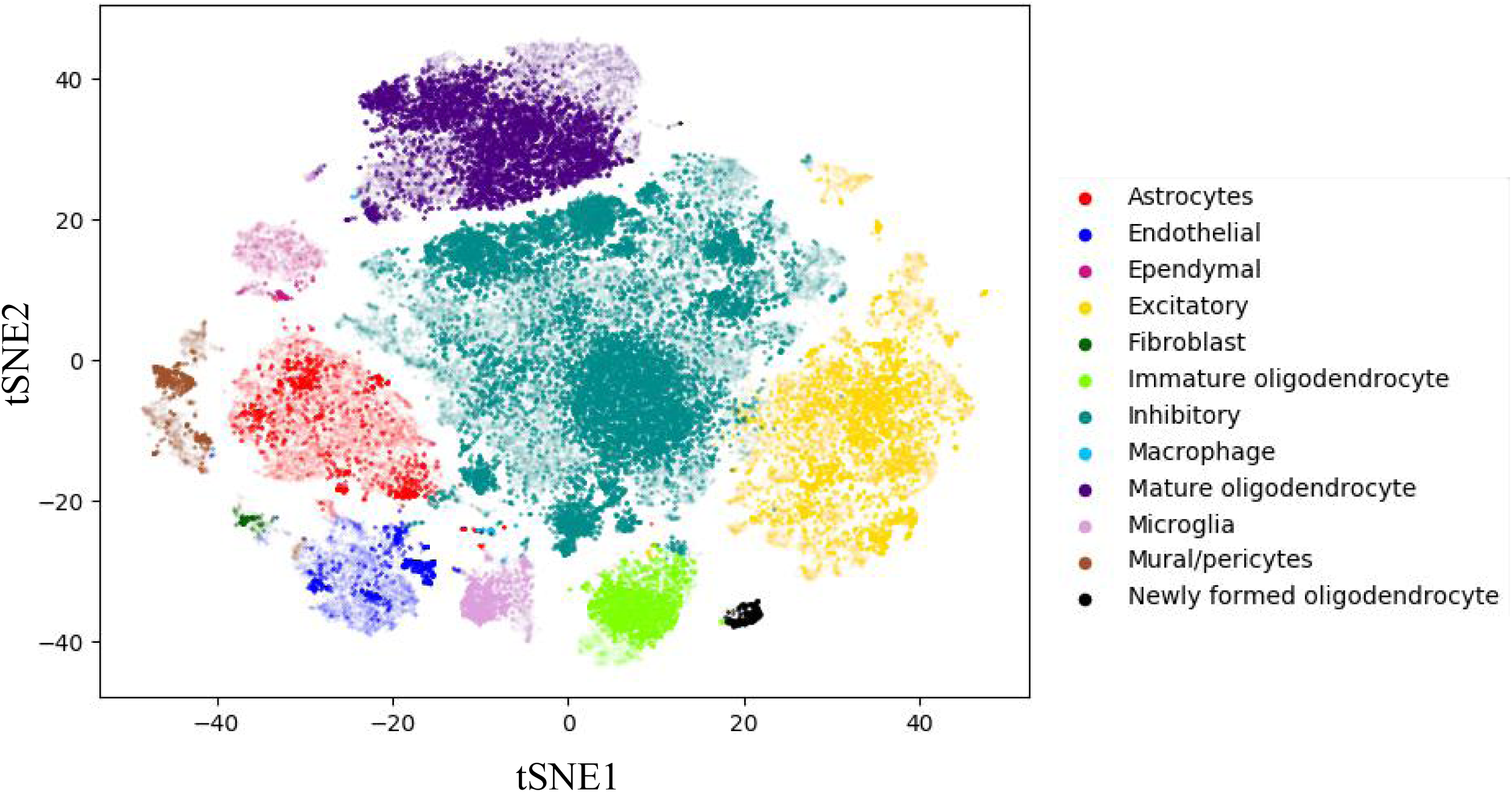
label transfer by inClust The t-SNE plot of the latent unified representation for scRNA-seq data and MERFISH data in inClust colored by predicted cell types. The normal part is from reference (scRNA-seq), and the blurred part is from target (MERFISH). The label is transferred from reference to target.

**Figure S3.**
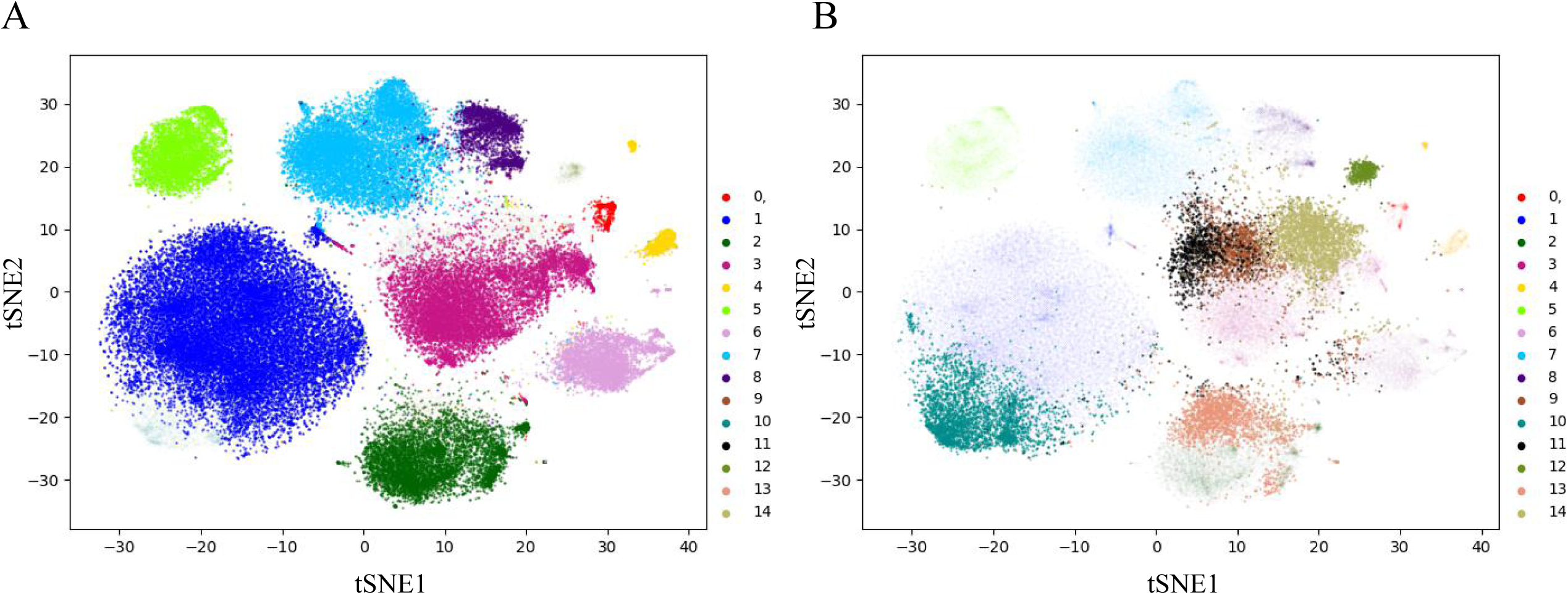
guided clustering by inClust A. The t-SNE plot of the latent unified representation for the heart scRNA-seq data in inClust is colored by cluster. The normal part corresponds to semi-supervised clusters, which could be labeled by reference. The blurred part corresponds to unsupervised clusters. B. The t-SNE plot of the latent unified representation for the heart scRNA-seq data in inClust is colored by clusters. The normal part corresponds to unsupervised clusters, which are annotated by marker genes. The blurred part corresponds to semi-supervised clusters.

**Figure S4.**
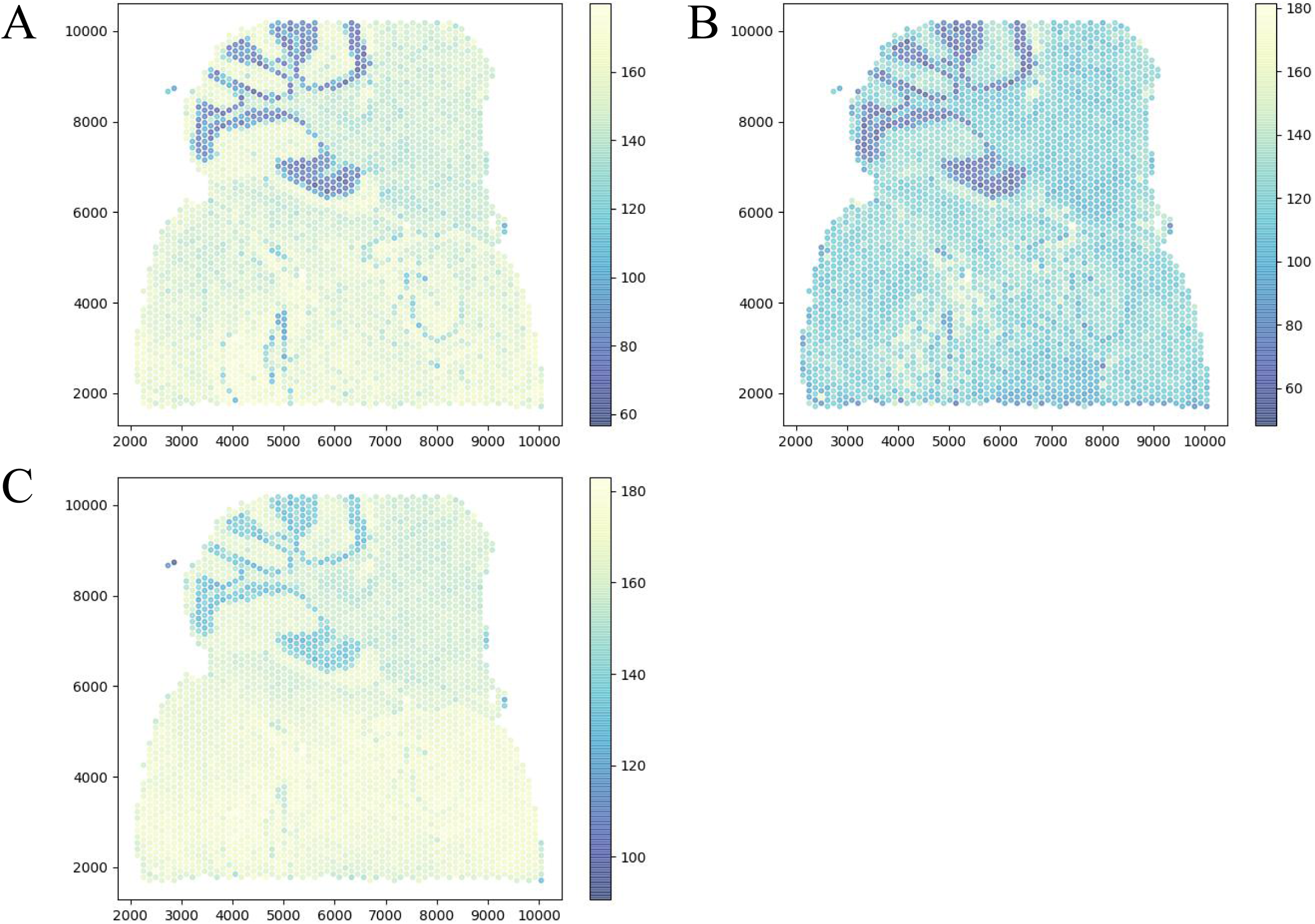
the smoothed RGB values of histological image A. B. C. the smoothed RGB values of histological image, respectively.

